# Extraction Optimization, Characterization, and Antioxidant Capacity of Phenolics from Cowpeas (*Vigna Unguiculata*)

**DOI:** 10.1101/2022.10.24.513571

**Authors:** Daniel M. Webber, Navam S. Hettiarachchy, Ronny Horax

**Affiliations:** Institution where work was performed: Department of Food Science, University of Arkansas, Fayetteville, Arkansas, USA

**Keywords:** Surface response, phenolic, cowpea, oxidative stability, DPPH, radical scavenging, optimization, *Vigna unguiculata*

## Abstract

Ethanol-water extraction of phenolics from cowpeas was modeled and optimized by response surface methodology (RSM). The ethanol concentration and extraction temperature were shown to have a significant effect on phenolic extraction and antioxidant capacity. Modeling predicted that extraction of phenolics from cowpea flour for 42.8 minutes at 58.6°C with 58.4% ethanol would maximize the radical scavenging capacity of solutes. Extraction of phenolics under these optimized conditions yielded 11.05 ± 0.10 mg chlorogenic acid equivalents (CAE)/g cowpea flour. These extracts contained 10.41 % ± 0.11% phenolics by weight and had an antioxidant capacity of 0.45 ± 0.02, closely approximating the predicted phenolic content of 10.11% ± 0.44% and antioxidant capacity of 0.42 ± 0.04. Extracted material was characterized by HPLC, and the predominant phenolic compounds detected were epicatechin and ferulic acid. Cowpea’s low cost, ease of storage, and high antioxidant capacity reflect their potential for use as a naturally-derived antioxidant additive in foods.

## INTRODUCTION

Lipid oxidation is a radical-mediated process that is often accelerated by light, temperature, or metal ions (1). Food quality attributes such as aroma, flavor, and color can be negatively affected by the oxidative process. Synthetic antioxidants such as tertiary butylhydroquinone (TBHQ), butylated hydroxytoluene (BHT), and butylated hydroxyanisole (BHA) are often added to foods to slow the rate of lipid oxidation. Although these compounds are effective, there is interest in replacing them with naturally-derived alternatives (2, 3). Spices such as rosemary, oregano, and sage have demonstrated strong antioxidant activity (4, 5). Furthermore, extracts from green tea, evening primrose seed, sage, and thyme have shown effectiveness at limiting oxidation in bulk oil systems (6–8). However, natural antioxidants are typically more expensive, yet inferior in quality compared to synthetic counterparts (9). Therefore, a need exists for an effective yet inexpensive natural antioxidant.

Cowpea is a drought-tolerant crop that improves the soil quality through nitrogen fixation. The seeds are a rich source of protein, vitamins, and minerals (10). These beneficial qualities have led to the propagation of cowpeas throughout the world (11). Previous studies have demonstrated that certain varieties of cowpea have high levels of phenolic compounds (12, 13). However, research has yet to optimize the extraction of cowpea phenolics or evaluate phenolic extracts as antioxidants. The solvent extraction of phenolics is dependent upon many factors including the solvent type, extraction temperature, solvent-to-solid ratio, extraction duration, particle size, and agitation (14). The principles of mass transfer are well established and their effects on phenolic extractions have been thoroughly explored (15–17). These principles provide some guidance in designing extractions; however, factors such as optimum solvent polarity, extraction duration, and temperature are greatly dependent upon the chemical characteristics of the target compound. Since the concentration and type of phenolic compounds vary among plant materials, there is a need to optimize extraction conditions for each plant matrix.

Phytochemical extractions can be optimized by varying one factor at a time; however, this process is very time-consuming and does not account for the interaction between extraction conditions. Since phenolic extractions are often significantly affected by factor interaction, the use of sequential optimization methods can lead to erroneous conditions for extraction. In contrast, optimization by response surface methodology (RSM) considers the interaction between multiple factors. This statistical method first introduced by Box and Wilson (18) also reduces the time required to determine optimum conditions. RSM has been used effectively in optimizing the extraction of phenolics from several plant sources including, wheat (19), milled berries (15), and grape pomace (14).

The high phenolic content of cowpeas as reported by Cai and others (12) provides rational for evaluating them as a source of natural antioxidants. With worldwide availability and low cost, cowpeas may offer an affordable alternative to synthetic food antioxidants. RSM offers an effective means of optimizing the solvent extraction of phytochemicals. Therefore, this study evaluates a food-grade solvent system and use RSM to determine the optimum conditions needed to maximize the antioxidant capacity of phenolic extracts. Optimization experiments will also be used to model the response of individual phenolics to the extraction conditions, and the optimized phenolic extract will be tested for radical scavenging capacity.

## MATERIALS AND METHODS

### Materials

Sodium carbonate, 2,2-Diphenyl-1-picrylhydrazyl (DPPH) and reagent-grade methanol, ethanol, petroleum ether, and ethyl acetate were obtained from VWR International (West Chester, Pa.). Reagents used in HPLC were of analytical grade and purchased from VWR International. Phenolic acid standards, Folin Ciocalteu’s reagent, and propylene glycol were from Sigma Chemical Co. (St. Louis, Mo.). Phenolic acids used in HPLC quantification included benzoic acid, caffeic acid, (±)-catechin, cinnamic acid, *o*-coumaric acid, *p*-coumaric acid, 2,4-dimethoxybenzoic acid, (-)-epicatechin, gallic acid, ferulic acid, gentisic acid, *p*-hydroxybenzoic acid, protocatechuic acid, sinapic acid, syringic acid, and vanillic acid. Tertiary butylhydroquinone (TBHQ) was obtained from Easton Chemical Co. (Kingsport, TN), and butylated hydroxy toluene (BHT) was purchased from Sigma-Aldrich.

### Sample preparation

Three cowpea varieties including Arkansas 95-104, Black Crowder, and Louisiana Purplehull were selected for use based on their high phenolic content, as reported by Cai (2003). Cowpea seeds were provided by Dr. Jalaluddin at the University of Arkansas, Pine Bluff; seeds were stored at 4°C until use. To facilitate phenolic extraction, seeds were ground to a powder by an IKA M20 universal mill (Wilmington, NC); the resulting material was passed through a 250μm sieve to obtain flour of uniform particle size. Sifted flour was tested for moisture, crude fat, and protein content by AACC methods: 44-15A, 30-26, 46-08, respectively.

### Selection of Extraction Conditions

Two preliminary steps were necessary before the application of response surface methodology (RSM). In the first step, a phenolic variety was selected for optimization based on phenolic yield from methanol extractions. Methanol served well as a solvent because of its good selectivity towards phenolic compounds and its common use in laboratory phenolic extractions. Solvent extraction was conducted in a temperature-regulated water bath. For each variety, 10g of cowpea flour was transferred into a 250 mL

Erlenmeyer flask containing 100 mL of 60°C methanol. Each flask was fitted with a water-cooled condenser and maintained at 60°C for 2 h with continuous stirring. Following extraction, samples were cooled to room temperature, centrifuged at 5,000 x g in a Beckman J2-21 centrifuge (Fullerton, CA), and vacuum filtered through a #4 Whatman filter; supernatants were rotary evaporated at 40°C until a 20mL volume remained. Concentrated samples were then freeze-dried, ground with a mortar and pestle, and stored at 4°C until use. A colorimetric method adapted from Singleton and Rossi (20) was used to determine the total phenolic content of each sample. The phenolic content was used to select a cowpea variety for further study and optimization.

The second preliminary step used sequential extractions to select the range of the factors used in response surface optimization. Since this step only assisted in the RSM design it was conducted without repetitions. The sequential optimization involved maintaining the treatment conditions for temperature and time while testing the level of extracted phenolics at various ethanol concentrations. The percent ethanol yielding the highest phenolic content was then fixed, temperature was maintained constant, and time was varied. Finally, percent ethanol and time were fixed at optimum values, and the effects of temperature were determined. This preliminary screening allowed for the effective selection of treatment levels for RSM design.

### Total Phenolic Content

The total phenolic content of freeze-dried cowpea extracts was determined by a colorimetric method adapted from Singleton and Rossi (20). This method involved solubilizing each sample in 1mL of 15% methanol followed by the addition of 5 mL of 0.2 N Folin-Ciocalteu reagent to each sample. Sample solutions were then vortexed thoroughly (10 sec) and incubated for 5 min before adding 4 mL of 708mM Na_2_CO_3_. After a final 10 sec of vortexing, samples were held in a 24°C water bath for 2 h before measuring absorbance at 765nm in a 1 cm cell using a Shimadzu UV-1601 spectrophotometer (Kyoto, Japan). Phenolic content was determined by use of a standard curve, and results were expressed in terms of chlorogenic acid equivalents (CAE). The term “phenolic yield” refers to quantity the phenolic compounds (mg CAE) extracted per gram cowpea flour. Phenolic content was defined as the percent phenolics detected in freeze-dried extract (w/w, CAE: extract) (21).

### Response surface optimization

A uniform precision, central composite design was used to evaluate the effects of temperature (T), treatment time (t), and percent ethanol (E) upon phenolic extraction and antioxidant capacity. The Experimental design included 5 replications at the model’s center point and 5 levels for each of the above independent variables, including the following factor levels: −1.68, −1,0, +1, +1.68 (Table S1). Each of the 19 experiments was performed in duplicate, and the coded variables (T_c_, t_c_, and E_c_) were determined as shown in Equation 1. Data from extractions were fitted to a response surface by the second-order polynomial displayed in Equation 2. The optimum extraction conditions for antioxidant capacity were determined with JMP 5.1 (SAS Institute Inc). Additionally, a surface response model was composed of total phenolic content serving as the independent variable rather than antioxidant capacity.

**Equation 1.** Coding of response surface variables

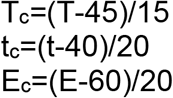

The variables T_c_, t_c_, and E_c_ represent the coded temperature, treatment time, and percent ethanol, respectively; and the variables T, t, and E represent the un-coded temperature, treatment time, and percent ethanol, respectively.

**Equation 2.** General response surface polynomial

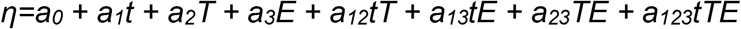

The term η represents the response variable, *a_0_* is a function representing the center point conditions, *a_1_, a_2_, a_3_* represent the principle effects associated with each factor, and *a_12_, a_13_, a_23_, a_123_* represent interactions between variables.

### Antioxidant Capacity

The antioxidant capacity of samples from RSM was determined by a DPPH^·^ (1,1-Diphenyl-2-picrylhydrazyl) radical scavenging method described by Sanchez et al. (22). Freeze-dried cowpea extract was dissolved in methanol to obtain concentrations of 10, 5, 2.5, 1, 0.5 mg extract/mL methanol. Because of its instability, a fresh solution of 25mg DPPH^·^L^-1^ methanol was prepared just before each use. A standard curve was established with DPPH^·^ concentration, and a volume of 0.1 mL of each sample was added to 3.9 mL of the DPPH^·^ methanol solution. Samples were vortexed for 10 seconds and incubated in the dark for 30 minutes. Absorbance was then measured at 515nm using a Shimadzu UV-1601 spectrophotometer. Sample absorbance was used to calculate EC_50_, which is defined as the concentration of sample per concentration DPPH^·^ required to decrease the steady-state concentration of DPPH^·^ by 50%. EC_50_ expresses the dose-dependent effects of antioxidant extract and was interpolated from a regression of percent DPPH^·^ remaining versus extract concentration divided by initial DPPH^·^. The inverse of EC_50_ is used in this paper to express antioxidant capacity.

### Model Verification

Following optimization by RSM, a final extract was prepared in triplicate using the optimum values for temperature, time, and percent ethanol. The validity of the optimization process was then evaluated by comparing the predicted antioxidant capacity from the RSM model with the actual antioxidant capacity obtained from extracts.

### HPLC Analysis of Phenolic Constituents

Phenolic compounds present in the optimized extract were identified and quantified by the HPLC method of Cai et al. (12) using 16 phenolic acid calibration standards, diluted to 5, 10, 20, and 50 μg/mL methanol. The optimized extract was prepared for analysis by dissolving 0.05g of the extract in 0.2 mL of methanol. The resulting solutions were then filtered through a 0.2-μm PVDF Target Syringe Filter (Natl. Scientific, Duluth, Ga., U.S.A.). After a standard curve was established, samples were analyzed on a Hewlett-Packard Liquid Chromatograph model 1090 (Agilent Technologies, Inc., Palo Alto, CA., USA) using a TSK-GEL Super-ODS (Supelco, Bellefonte, PA, USA) column for phenolic acid separation. The mobile phase consisted of solvent A (HPLC grade water with 0.1% trifluoroacetic acid) and solvent B (acetonitrile with 0.1% trifluoroacetic acid). Solvent flow rate and temperature were maintained at a respective 1.0 mL/min and 37 °C throughout the analysis. The initial solvent was composed of 100% solvent A; upon injection of 4 μL of the analyte, solvent B was increased linearly from 0% - 10% within 7 minutes. The solvent composition was then maintained isocratically for 3 minutes followed by a linear increase in solvent B from 10% to 40% during a 20 min period. The solvent composition was maintained at this level for 2 min, and solvent B was then returned to 0% within 3 min. HPLC grade methanol, 100%, was used to wash the column before and after sample analysis. The absorbance of eluting compounds was monitored by UV detector at 254 nm and phenolic constituents were quantified by comparison to the linear regressions of standard compounds.

## RESULTS AND DISCUSSION

### Variety Selection

Methanol extractions from Black Crowder, Arkansas 95-104, and Louisiana Purplehull yielded 2.57 ± .05, 0.77 ± .05, and 1.01 ± .08 mg CAE/g flour, respectively. ANOVA testing indicated that the Black Crowder cowpea variety contained higher levels of extractable phenolic acids compared to Arkansas 95-104 and Louisiana Purplehull; therefore, Black Crowder was selected for further optimizations.

### Response Surface Optimization

Response surface methodology was used to optimize the ratio of ethanol to water, extraction time, and temperature. Ethanol-water was used for extraction based on the solubility profile of phenolic compounds. The presence of water in the solvent system has been reported to accelerate mass transfer by swelling the solid matrix, which allows better solvent access. Extraction time and temperature were also selected for optimization based on prior work with phenolic acid extraction (15, 19, 24).

### Phenolic Content

The RSM regression fitted well to the experimental results for phenolic content (r^2^ = 0.97). Table S2 shows that the phenolic content of extracts was significantly affected by temperature (p<0.0001), solvent ratio (p<0.0001), and the interaction between temperature and solvent ratio (p=0.0121). Extraction time did not have a significant effect on the model. As Figures 1 and 2 indicate, the level of extracted phenolics increased linearly with increasing temperature. The effects of temperature on extract concentration can be attributed to an increase in solubility and diffusion coefficient (15, 24). The linear relationship between temperature and concentration placed the optimum temperature outside of the experimental range; this linearity indicated that the temperature did not reach a level where degradation outpaced phenolic release. The interaction between solvent ratio and temperature created a rising ridge that at 40 min migrates diagonally from 44% ethanol at 20°C towards 58% ethanol at 70°C (Figure 2). The diagonal shift in the response variable has been reported in the case of anthocyanin extraction from black currants (15). The interaction between the optimum solvent ratio and temperature is related to the activity coefficient of the phenolic compounds, which is temperature dependent (15, 25).

**Figure 1.**
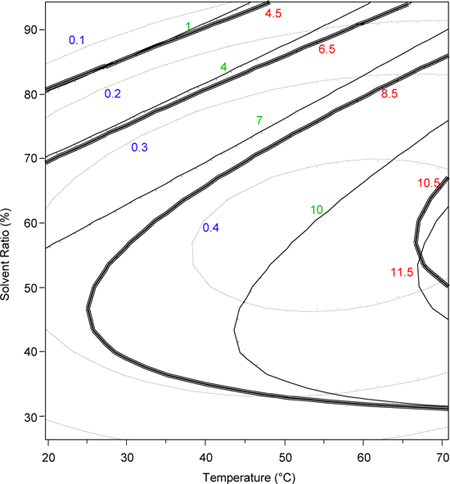
Effect of solvent concentration and temperature on phenolic content, phenolic yield, and antioxidant capacity of extracts, measured after 40-minutes. The contour plot displaying phenolic content (solid dark line, percent of dry weight), phenolic yield (single solid line, mg CAE / g cowpea flour), and antioxidant capacity (single broken line, 1/EC_50_).

**Figure 2.**
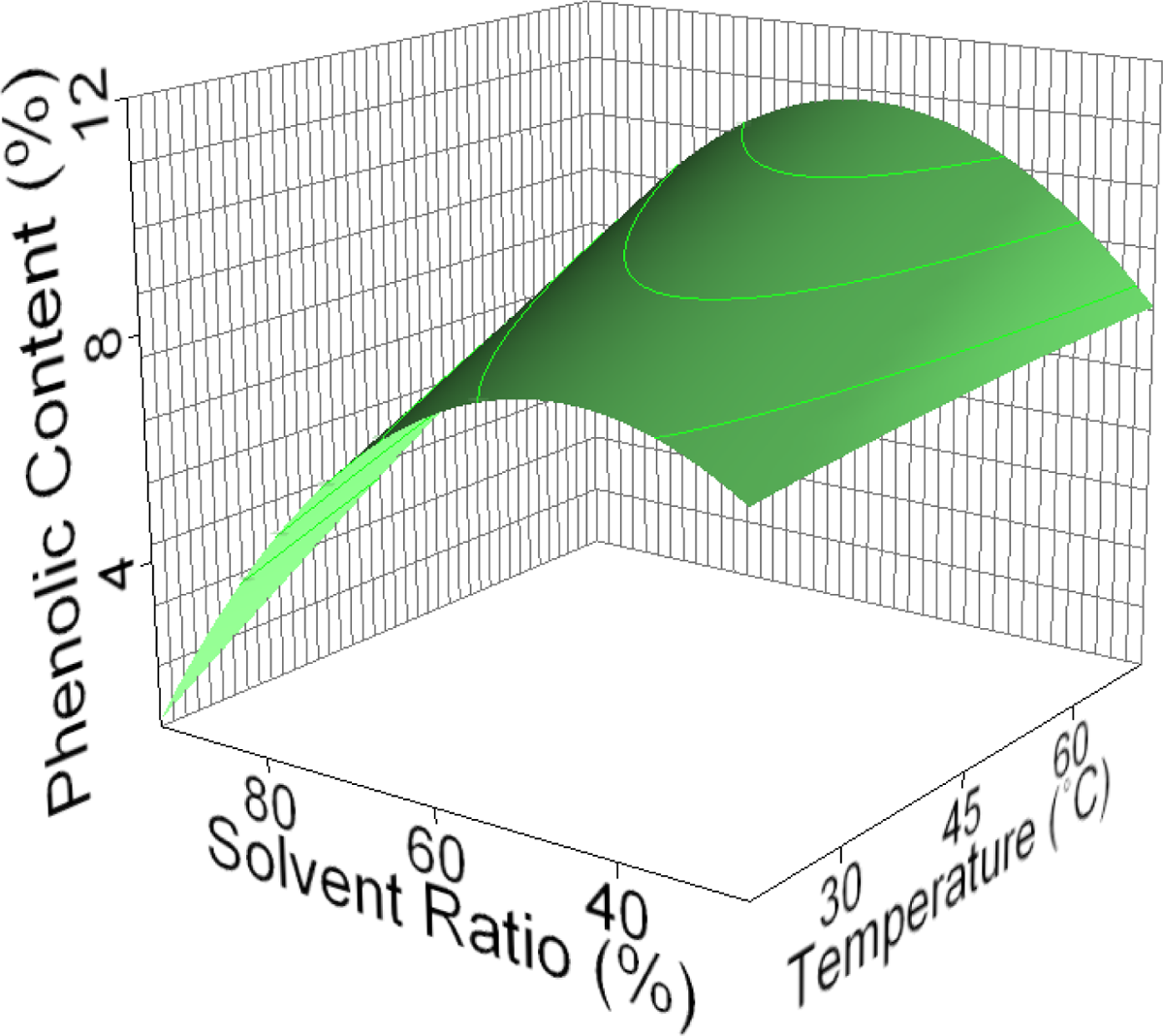
Effect of solvent concentration and temperature on phenolic content of extracts, measured after 40-minutes. Phenolic content of extracts is expressed as a percent of dry weight (w/w). Solvent ratio refers to the concentration of ethanol in water.

**Figure 3.**
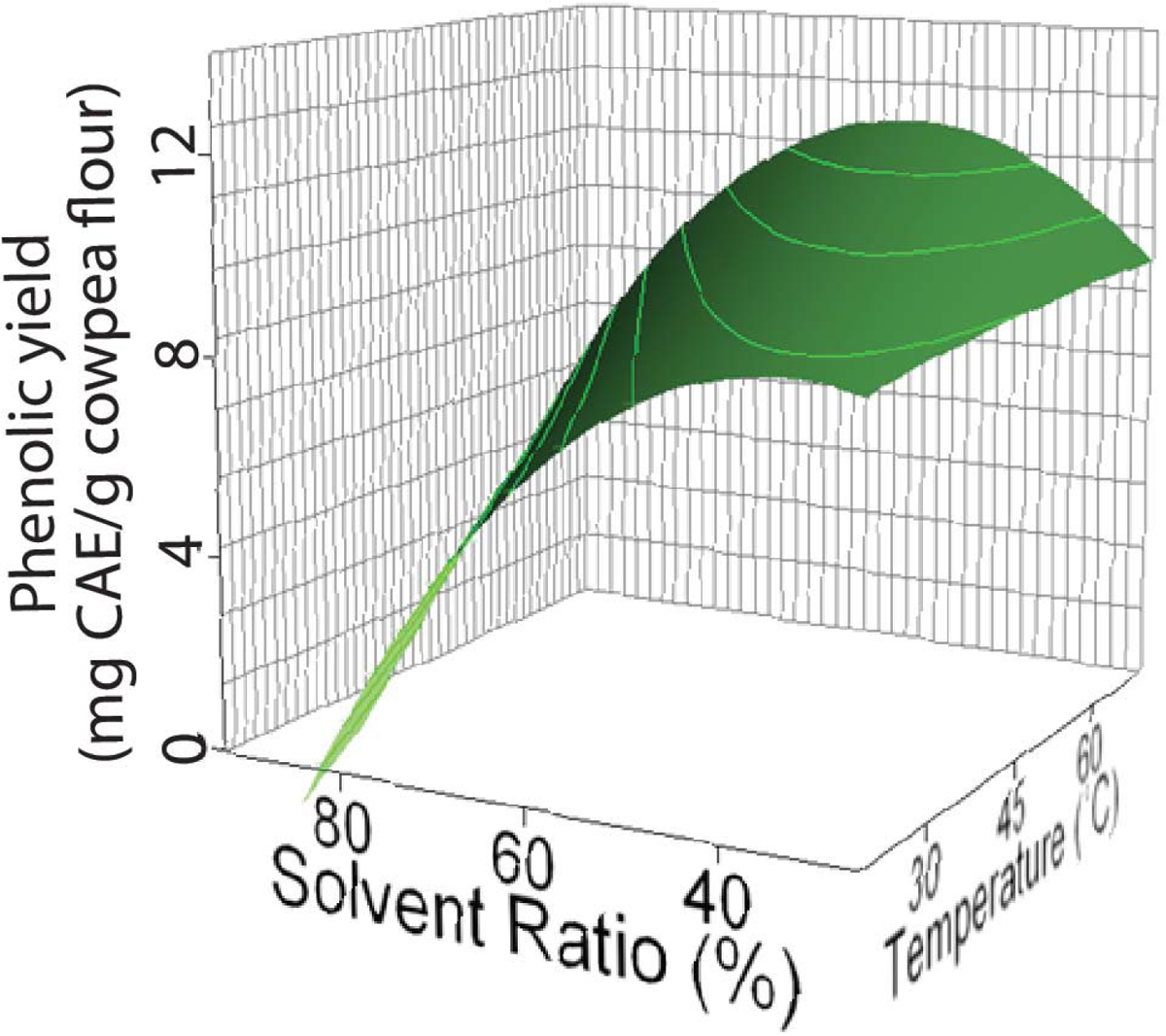
Effect of solvent ratio and temperature on phenolic yield, measure after 40 minutes. Phenolic yield is expressed in chlorogenic acid equivalents (CAE)/g cowpea flour. The concentration of ethanol in water (v/v) is denoted as solvent ratio (%).

### Phenolic Yield

The predictive equation used to describe phenolic yield was highly correlated (r^2^=0.99) with experimental data. Statistical analysis, as shown in Table S2, indicates that yield was significantly affected by temperature, solvent ratio, and the interaction between temperature and solvent ratio. The optimum extraction conditions within the experimental bounds were predicted at 70.2°C, 73.6 minutes, and 53.6% ethanol.

### Antioxidant Capacity

The data from optimization experiments were used to create a predictive model for antioxidant capacity. A secondary polynomial was found to best represent the data, and Table S3 displays the coefficients from the predictive equation. Regression analysis, as shown in table S2, indicates that the RSM model adequately describes the data with an r^2^ of 0.90. Antiradical capacity was significantly affected by the extraction temperature (p=0.0207) and solvent ratio (p=0.0168); however, extraction time did not have a significant effect on the among of phenolic that were extracted. As Figures 1 and 4 indicate, antioxidant capacity increased with increasing temperature, reached an optimum, then decreased with further increases in temperature. Solvent ratio also had a similar effect on antioxidant capacity, producing an optimum point near 60% under most conditions. Previous research indicates that the optimum ethanol concentration for antioxidant extraction is dependent upon the plant material. However, ethanol-water extractions from wheat (19), borage seeds (26), and evening primrose meal (27) have demonstrated the general effectiveness of a 50-60% ethanol solvent in extracting antioxidant compounds from plant material. Figure 1 demonstrates that the response surface for antioxidant capacity was similar to the response profile for phenolic content. Further analysis indicated that phenolic content and antioxidant capacity were positively correlated (r^2^ of 0.78, Figure 5), consistent with prior reports of antioxidative activity of phenolic compounds (28).

**Figure 4.**
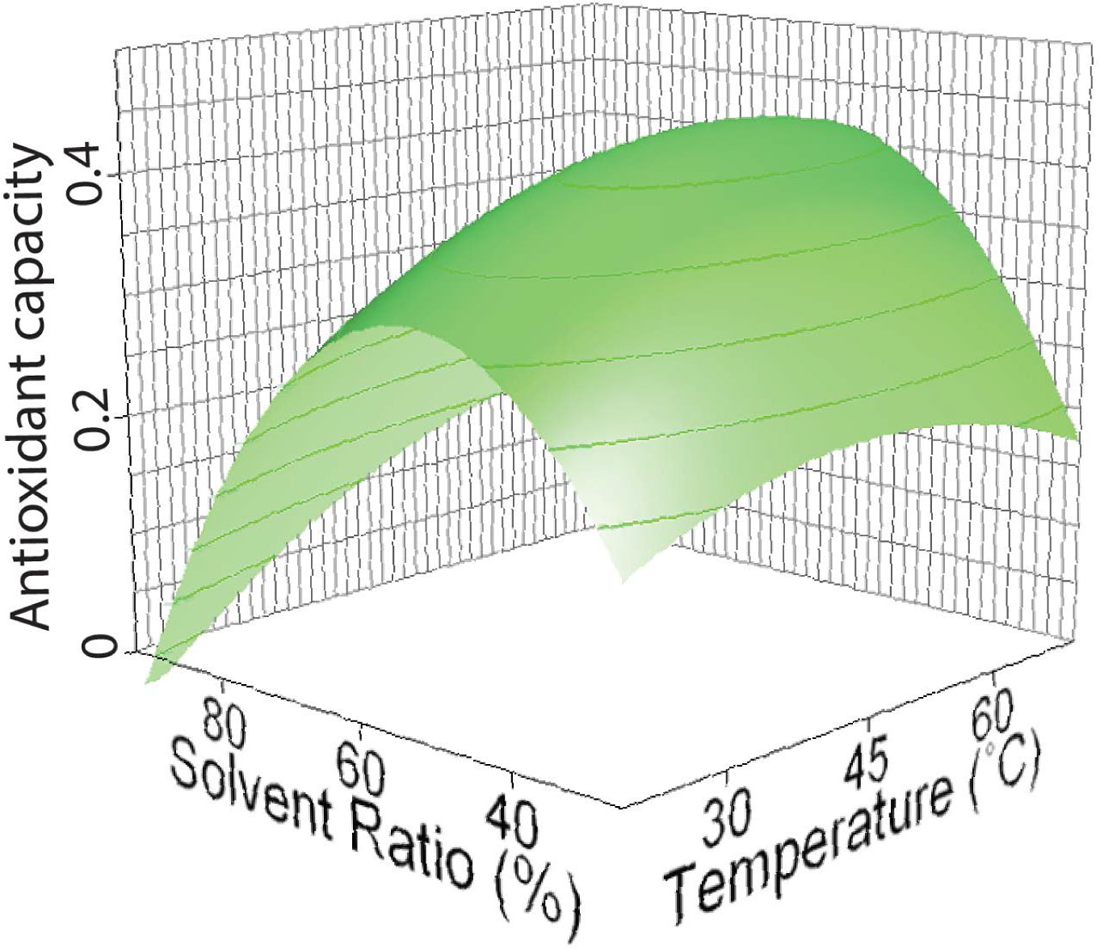
Effect of solvent ratio and temperature on antioxidant capacity of extracts, measure after 40 minutes. Antioxidant capacity is reported here as the inverse of EC50 for a sample, where EC50 represents the ratio of sample-to-DPPH^·^ needed to decrease the steady-state concentration of DPPH^·^ by 50%. The concentration of ethanol in water (v/v) is denoted as solvent ratio (%).

**Figure 5.**
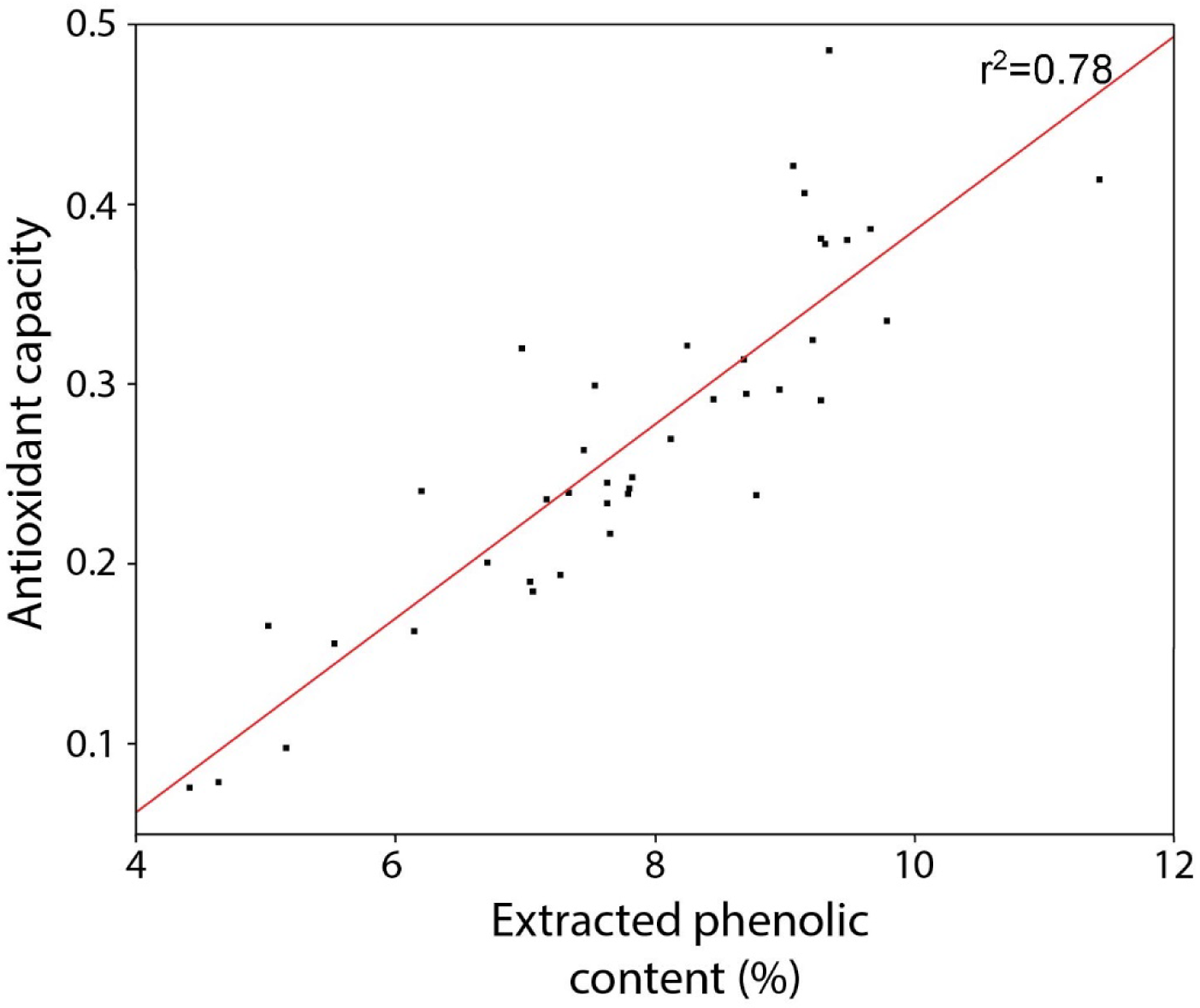
Correlation between phenolic content (i.e. purity) and antioxidant capacity of samples that were extracted under various conditions. Phenolic content of extracts is expressed as a percent of dry weight.

### Optimized Extraction

The optimum conditions for maximizing antioxidant capacity were 58.6°C, 42.8 min, and 58.4% ethanol. By substituting the optimized conditions into the response surface models, values for phenolic content, phenolic yield, and antioxidant capacity were predicted (Figure 6) as 10.11% ± 0.44%, 10.66 ± 0.45 mg CAE/g cowpea flour, and 0.43 ± 0.04, respectively. The predicted response was validated using a final extraction, carried out under the optimized conditions for antioxidant capacity (58.6°C, 42.8 min, and 58.4% ethanol). Extracts from this reaction had a phenolic content (i.e. purity) of 10.41% ± 0.11% and an antioxidant capacity of 0.45 ± 0.02. The yield of this optimized reaction was 11.05 ± 0.10 mg CAE/g cowpea flour. The actual phenolic content, phenolic yield, and antioxidant capacity of extracts from the optimized reaction were highly similar to the predicted values.

**Figure 6.**
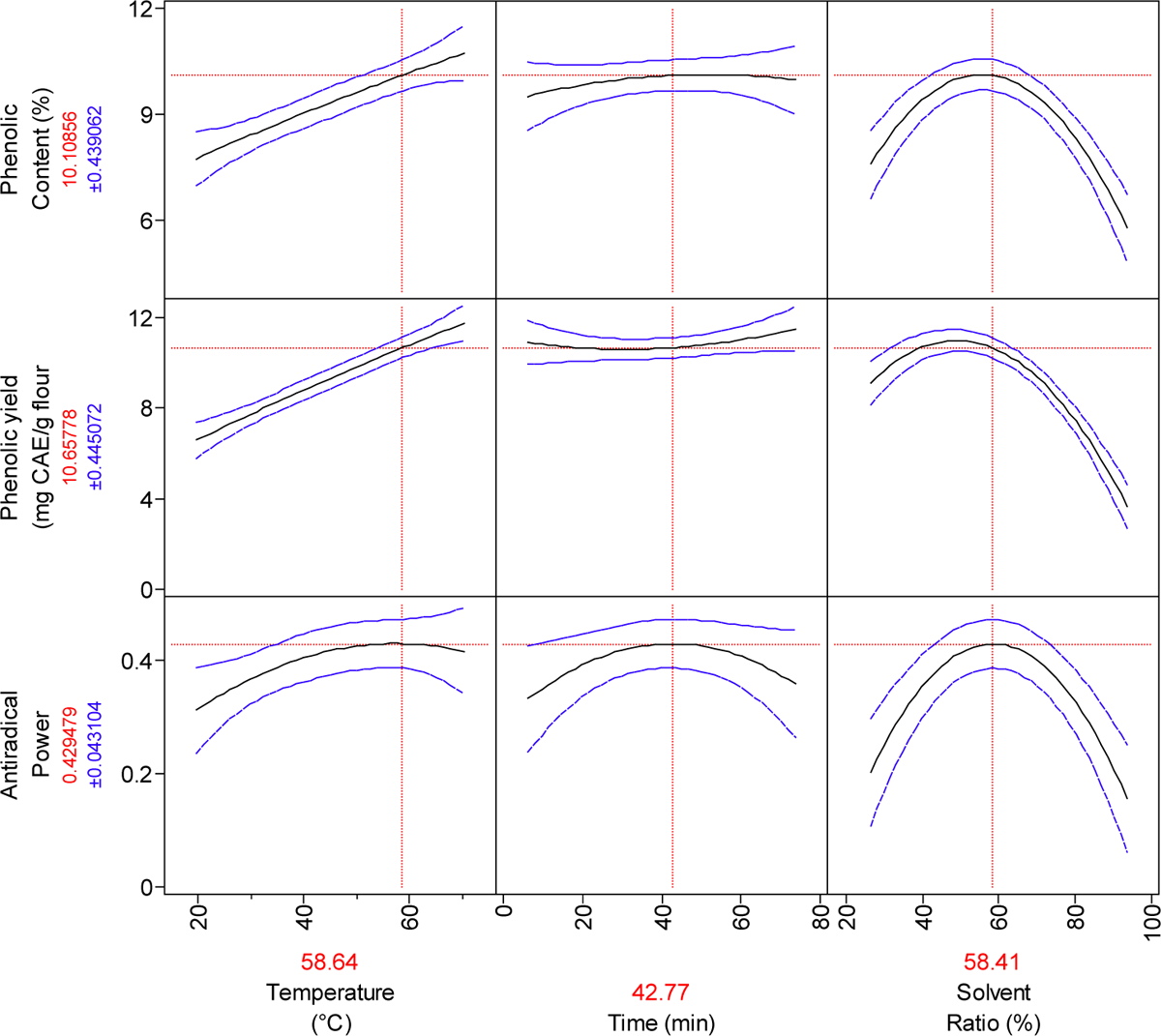
Predicted effect of solvent temperature, incubation time, and solvent concentration (ethanol/water, v/v) on phenolic content (%), phenolic yield, and antioxidant capacity of extracts. Phenolic content of extracts is expressed as a percent of dry weight (w/w). Phenolic yield is expressed in chlorogenic acid equivalents (CAE)/g cowpea flour. The predicted responses ± SD are shown in red text on the Y-axis. Optimized extraction conditions are show in red are shown in red text on the X-axis.

**Figure 6.**
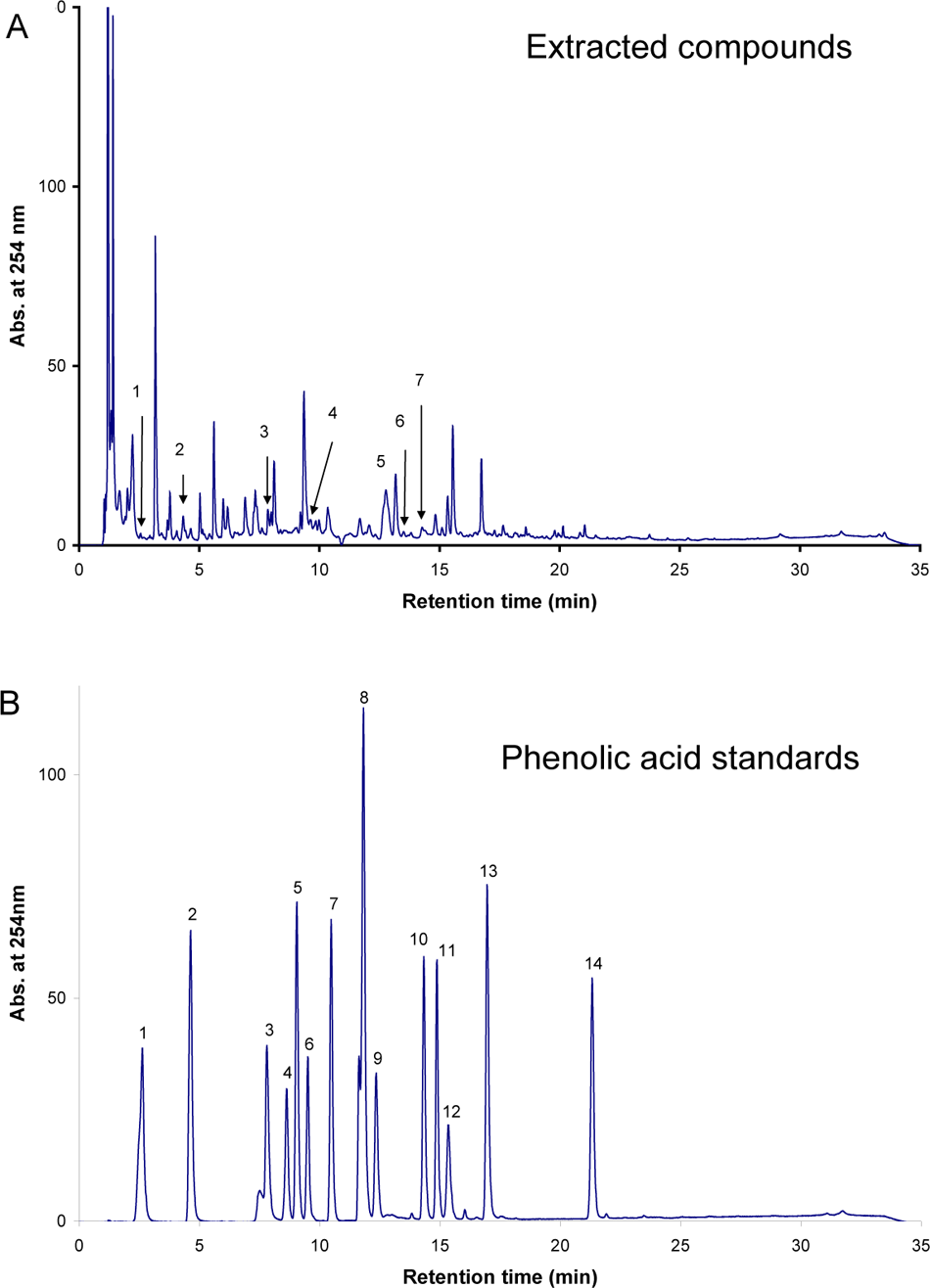
HPLC Chromatogram. **a)** Chromatogram from optimized extract. The numbers above peaks are for phenolic compound identification as follows: 1, gallic acid; 2, protocatechuic acid; 3, vanillic acid; 4, (-)-epicatechin; 5, *trans*-ferulic acid; 6, sinapic acid; 7, *p*-Hydroxybenzoic acid. **b)** Chromatogram of phenolic standards. The numbers above each peak are for phenolic comopund identification as follows: 1, gallic acid; 2, protocatechuic acid; 3, gentistic acid; 4, catechin; 5, vanillic acid; 6, caffeic acid; 7, syringic acid; 8, (-)-epicatechin; 9, p-courmaric acid; 10, *trans*-ferulic acid; 11, sinapic acid; 12, *p*-hydroxybenzoic acid; 13, o-coumaric acid; 14, *trans*-cinamic acid

### Phenolic Constituents

We next sought to quantify phenolic acid compounds that were extracted under optimized conditions. Seven phenolic acid compounds were detected (table 1) in the ethanol extracts by performing HPLC and comparing retention times to flavan-3-ol standards (figure 6). The most abundant phenolic compound present in the cowpea extract was (-)-epicatechin (table 1), which is a flavan-3-ol commonly found in black grapes, red wine, and green tea (29). The second most abundant phenolic constituent was *trans*-ferulic acid, which is a hydroxycinnamic acid that is found throughout the plant world (30). Sinapic acid was also detected in the extract; and several hydroxybenzoic acids were quantified, including benzoic acid, gallic acid, protocatechuic acid, and vanillic acid.

**Table 1.** Phenolic content and yield of optimized extract.

| Compound | Phenolic content<br>(mg/100 g extract) |
| --- | --- |
| gallic acid | 7.80 |
| protocatechuic acid | 13.62 |
| vanillic acid | 11.89 |
| (-)-epicatechin | 222.78 |
| *trans*-ferulic acid | 82.29 |
| sinapic acid | 14.83 |
| *p*-hydroxybenzoic acid | 20.55 |

## Conclusions

This study demonstrated (i) that cowpeas contain significant amounts of phenolic compounds, including (-)-epicatechin and trans-ferulic acid, (ii) phenolic acids in cowpeas are effective antioxidants, and (iii) the level of antioxidant capacity in cowpea extracts can be predicted accurately by modeling the effects of temperature and solvent concentration on these factors.

