## Supplemental Table 1 for "Extraction Optimization, Characterization, and Antioxidant Capacity of Phenolics from Cowpeas (*Vigna Unguiculata*)"

SUPPLEMENTAL RESULTS

**Table S1**. Conditions for RSM experiments

| Expt. | $T_c$ (°C) | $t_c$ (min) | $E_c$ (%) | T (°C) | t (min) | E (%) |
|---|---|---|---|---|---|---|
| 1 | -1.68 | 0 | 0 | 19.8 | 40 | 60 |
| 2 | -1 | -1 | -1 | 30 | 20 | 40 |
| 3 | -1 | -1 | +1 | 30 | 20 | 80 |
| 4 | -1 | +1 | -1 | 30 | 60 | 40 |
| 5 | -1 | +1 | +1 | 30 | 60 | 80 |
| 6 | 0 | -1.68 | 0 | 45 | 6.4 | 60 |
| 7 | 0 | 0 | -1.68 | 45 | 40 | 26.4 |
| 8 | 0 | 0 | 0 | 45 | 40 | 60 |
| 9 | 0 | 0 | 0 | 45 | 40 | 60 |
| 10 | 0 | 0 | 0 | 45 | 40 | 60 |
| 11 | 0 | 0 | 0 | 45 | 40 | 60 |
| 12 | 0 | 0 | 0 | 45 | 40 | 60 |
| 13 | 0 | 0 | 0 | 45 | 40 | 60 |
| 14 | 0 | 0 | +1.68 | 45 | 40 | 93.6 |
| 15 | 0 | +1.68 | 0 | 45 | 73.6 | 60 |
| 16 | +1 | -1 | -1 | 60 | 20 | 40 |
| 17 | +1 | -1 | +1 | 60 | 20 | 80 |
| 18 | +1 | +1 | -1 | 60 | 60 | 40 |
| 19 | +1 | +1 | +1 | 60 | 60 | 80 |
| 20 | +1.68 | 0 | 0 | 70.2 | 40 | 60 |

[a]T, t, and E represent treatment temperature, time, and ethanol concentration, respectively; and Tc, tc, and Ec represent coded variables
