## Supplemental Table 2 for "Extraction Optimization, Characterization, and Antioxidant Capacity of Phenolics from Cowpeas (*Vigna Unguiculata*)"

SUPPLEMENTAL RESULTS

**Table S2.** Significance of beta values from

| | Response variable | | |
|---|---|---|---|
| factor | Phenolic content[a] | Phenolic yield[b] | Antioxidant capacity[c] |
| temp | <0.0001 | <0.0001 | 0.0207 |
| time | 0.0609 | 0.2272 | 0.3917 |
| solvent ratio | <0.0001 | <0.0001 | 0.0168 |
| temp*time | 0.5169 | 0.8335 | 0.9615 |
| solvent ratio*temp | 0.0121 | 0.0003 | 0.2576 |
| time*solvent ratio | 0.1301 | 0.9565 | 0.1322 |

[a]Values are expressed in chlorogenic acid equivalents (CAE). Phenolic content equals mg CAE / 100 mg extract (dry weight basis). [b]Phenolic yield expressed as mg CAE / g cowpea flour (dry weight basis). [c]Antioxidant capacity equals $1/EC_{50}$.
