## Supplemental Table 3 for "Extraction Optimization, Characterization, and Antioxidant Capacity of Phenolics from Cowpeas (*Vigna Unguiculata*)"

SUPPLEMENTAL RESULTS

**Table S3.** Variable coefficients from polynomial equation describing antioxidant capacity

| | Temp | Time | Solvent Ratio | Antioxidant Capacity |
|---|---|---|---|---|
| Temperature | $-7.5x10^{-05}$ | $-2.5x10^{-06}$ | $6.1x10^{-05}$ | $2.1x10^{-03}$ |
| Time | . | $-7.0x10^{-05}$ | $6.3x10^{-05}$ | $5.2x10^{-04}$ |
| Solvent Ratio | . | . | $-2.2x10^{-04}$ | $-1.7x10^{-03}$ |
